# Blocking protein quality control degradation leads to structural stabilization of DHFR indel variants

**DOI:** 10.64898/2025.12.09.693117

**Authors:** Sven Larsen-Ledet, Caroline H. Suhr, Sarah Gersing, Celeste M. Hackney, Amelie Stein, Kaare Teilum, Rasmus Hartmann-Petersen

**Affiliations:** Department of Biology, University of Copenhagen, Ole Maaløes Vej 5, DK-2200 Copenhagen, Denmark

**Keywords:** folding, misfolding, protein stability, gene variants, proteostasis, PQC, UPS, ubiquitin, proteasome, chaperone, E3, yeast

## Abstract

Gene variants leading to insertions or deletions of amino acid residues (indels) often have detrimental consequences for the folding of the encoded protein. Yet at some positions indels are tolerated or result merely in partial unfolding and hypomorphic phenotypes. Typically unfolded proteins are targeted for protein quality control (PQC) degradation via the ubiquitin-proteasome system, which in yeast is mediated by specific E3 ubiquitin-protein ligases, including Ubr1 and San1. Here we systematically probed the folding of a library of indel variants in the DHFR protein using a sensitive yeast-based protein folding reporter. We show that deletion of Ubr1 or San1 leads to a greater fraction of folded DHFR indel variants, primarily positioned towards the N- and C-terminal regions in DHFR. Intriguingly, most of the DHFR indels that are structurally stabilized in the E3 knockout strains, are also stabilized at lowered temperatures and upon binding the DHFR inhibitor methotrexate. This suggests that blocking PQC degradation can restore function to partially unfolded hypomorph variants, thus providing a potential therapeutic avenue for protein misfolding diseases.

## Introduction

Dictated by their amino acid sequences, most proteins inherently fold into thermodynamically stable native conformations (1). However, stress conditions, such as elevated temperatures, the absence of essential cofactors or binding partners, or mutations that alter the amino acid sequence, can destabilize the native fold (2, 3). When this occurs, an increased fraction of the protein population will exist in unfolded or partially unfolded states. These non-native conformations often expose hydrophobic regions that are usually buried within the protein’s core, making them prone to participate in aberrant interactions with other cellular proteins (3, 4). Thus, by potentially disrupting normal protein interaction networks and contributing to proteotoxic stress, misfolded or unfolded proteins pose a significant threat to cells.

To mitigate this, cells have evolved a sophisticated protein quality control (PQC) system (2, 5, 6). Molecular chaperones recognize exposed hydrophobic regions in non-native proteins and catalyze folding to the native conformation (4, 7). In parallel, misfolded proteins also become targets for intracellular degradation (5). Central to this process are a group of E3 ubiquitin-protein ligases, including the yeast Ubr1 and San1, that display partially overlapping substrate specificities (8–13). These enzymes specifically recognize non-native proteins, by detecting exposed hydrophobic regions (14–16) and then catalyze conjugation to ubiquitin, which in turn marks the proteins for proteasomal degradation. This targeted degradation prevents the accumulation of toxic protein aggregates and helps maintain proteostasis. Thus, together, molecular chaperones and PQC E3s form a critical surveillance system that limits accumulation of non-native proteins.

Importantly, studies have shown that the PQC system is highly sensitive and does not exclusively select completely misfolded or non-functional proteins but may also recognize partially unfolded proteins that are still functional (8, 17, 18). Accordingly, PQC degradation can be a double-edged sword: while PQC-mediated clearance prevents the accumulation of potentially harmful protein species, it may also eliminate partially active proteins that could otherwise contribute to some vital cellular function. Consequently, understanding the mechanisms and specificity of the PQC system is critical. In particular, since this potentially opens therapeutic avenues for treating protein misfolding diseases (19–21). By preventing overly aggressive PQC degradation of structurally destabilized, but functional proteins, it may be possible to restore sufficient protein function to alleviate certain genetic diseases.

In this study, we systematically investigated the folding of a library of indel variants in the dihydrofolate reductase (DHFR) protein using a protein folding sensor. We show that deletion of the E3 ubiquitin ligases Ubr1 and San1 results in a higher proportion of folded DHFR indel variants. Notably, many of the DHFR indels that are structurally stabilized in the E3 knockout strains also show increased stability at lower temperatures and in the presence of the DHFR inhibitor methotrexate. These findings suggest that inhibiting PQC-degradation can help restore function to partially unfolded indel variants, highlighting a potential therapeutic strategy for treating protein misfolding disorders.

## Results

### Systematic assessment of PQC E3s on target protein folding in vivo

We previously applied our protein folding sensor, which is based on circular permutated orotate phosphoribosyltransferase (CPOP), to systematically assess the folding and stability of amino acid insertions and deletions (indels) in human DHFR in yeast cells (22). In this study, we extend that work by investigating how components of the protein quality control (PQC) system, specifically two E3 ubiquitin ligases, affect the folding and stability of DHFR indels *in vivo*. Ubr1 and San1 are two prominent PQC E3 ligases in yeast that target proteins for proteasomal degradation, functioning primarily in the cytosol and nucleus, respectively.

In the CPOP system, the *E. coli* orotate phosphoribosyltransferase (OPRTase) is circularly permuted so that the original N-terminal segment (residues 1–92) is relocated to the C terminus at position 213. The resulting CPOP enzyme remains active but is less structurally stable (23), and thus is not expected to significantly affect the stability of a protein inserted into the linker that joins the original C and N termini. However, even small structural alterations in the inserted protein are expected to disrupt CPOP folding, reducing OPRTase activity below the level needed to rescue a *ura5*Δ*ura10*Δ double deletion mutant yeast strain. Starting from the *ura5*Δ*ura10*Δ strain (hereafter referred to as WT strain) (22), we constructed a *ura5*Δ*ura10*Δ*ubr1*Δ triple mutant strain (hereafter referred to as *ubr1*Δ strain) and a *ura5*Δ*ura10*Δ*san1*Δ triple mutant strain (hereafter referred to as *san1*Δ strain). Each strain was transformed with the tiled DHFR indel library inserted into the CPOP linker as described before (22). Briefly, the DHFR indel libraries included sequences encoding: wild-type DHFR; a synonymous (silent) variant at each position; a nonsense (stop codon) mutation at each position; a single glycine insertion at each position; and a single amino acid deletion at each position. Following transformation, cells were plated on solid medium without uracil and incubated at both 30 °C and 35 °C for two days to select for variants adept at protein folding (**Fig. 1A**). After selection, plasmids were isolated and amplicons were prepared for next-generation sequencing (**Fig. 1A**). The resulting reads were processed using our in-house pipeline (22) to calculate a CPOP folding score for each indel variant relative to the synonymous variants in the WT, *ubr1*Δ, and *san1*Δ strains at both temperatures. The CPOP folding scores were rescaled such that a score of 0 represents the folding of stop codon variants, and a score of 1 represents the folding of synonymous wild-type variants. The datasets for the WT, *ubr1*Δ, and *san1*Δ strains include a CPOP folding score for 174 out of 177 (98%) possible synonymous wild-type variants, 185 out of 186 (99%) stop codon variants, 186 out of 187 (99%) insertion variants, and 186 out of 187 (99%) deletion variants. All data are included in the supplementary file (**Supplementary File 1**). Furthermore, sequencing read counts showed a high degree of correlation across the three replicates (**Supplementary Fig. 1-3**).

**Fig. 1.**
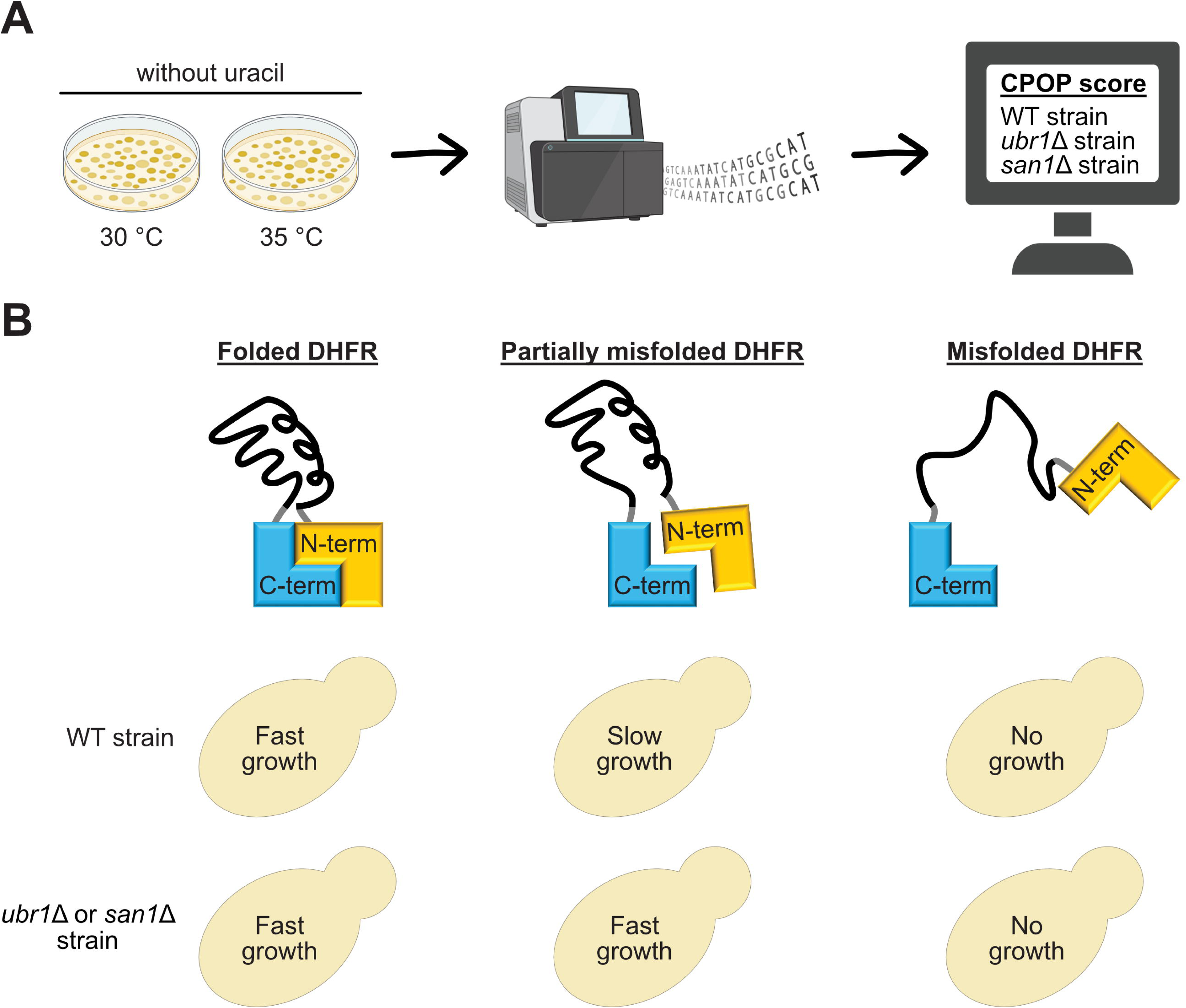
Assessing the impact of PQC E3 ligases on protein folding using CPOP. (A) Schematic overview of the experimental setup used to screen DHFR indel libraries in the WT, *ubr1*Δ, and *san1*Δ strains. Following transformation, cells were plated on medium without uracil and incubated at 30 °C and 35 °C for two days. After selection, cells were harvested for plasmid isolation and amplicon preparation, followed by next-generation sequencing. Sequencing reads were then processed to compute a CPOP folding score for each indel variant in the WT, *ubr1*Δ, and *san1*Δ strains at 30 °C and 35 °C. (B) The CPOP folding sensor consists of an N-terminal (yellow) and a C-terminal (blue) region that fold into the functional CPOP enzyme. This enzyme restores uracil prototrophy in the *ura5*Δ*ura10*Δ strain, which lacks endogenous OPRTase activity. Wild-type or indel variants of human DHFR (black) were inserted into the short linker region (grey) between the N- and C-terminal regions. The CPOP-DHFR constructs were transformed into three strains: the *ura5*Δ*ura10*Δ strain (WT strain) as well as a *ura5*Δ*ura10*Δ*ubr1*Δ (*ubr1*Δ strain) and *ura5*Δ*ura10*Δ*san1*Δ (*san1*Δ strain). If a DHFR indel variant is stable and well-folded (left), CPOP can fold and form a functional enzyme, enabling growth on medium without uracil in all three strains. A partially misfolded variant (middle) that is a substrate of Ubr1 or San1 shows slow growth in the WT strain due to proteasomal degradation, but fast growth in the *ubr1*Δ and *san1*Δ strains. In contrast, a fully misfolded variant (right) results in a complete growth defect across all three strains due to global and irreversible misfolding. Parts of the figure were created in BioRender. Hartmann-Petersen, R. (2025) https://BioRender.com/idshvgv.

In our assay, a well-folded DHFR indel variant is expected to support growth in the WT, *ubr1*Δ, and *san1*Δ strains (**Fig. 1B**). In contrast, a partially misfolded variant targeted for degradation by Ubr1 or San1 should display slow growth in the WT strain, but faster growth in the *ubr1*Δ and *san1*Δ strains (**Fig. 1B**). Finally, a completely misfolded variant, regardless of whether it is a substrate of Ubr1 or San1, is expected to exhibit a strong growth defect in all three strains (**Fig. 1B**). Therefore, we envision that the CPOP folding sensor is particularly useful for identifying hypomorphic indel variants that can refold and become functional when not targeted for degradation by Ubr1 or San1. In contrast, the sensor is not suited for identifying fully misfolded, amorphic variants that are substrates of Ubr1 or San1, as these variants show a similar growth defect regardless of the presence or absence of the E3 ligases.

### Deletion of PQC E3s leads to increased folding of DHFR indel variants

As we have shown previously (22), many DHFR indel variants were poorly folded and exhibited low CPOP folding scores in the WT strain at both temperatures (**Fig. 2**). Poor folding (CPOP folding score < 0.25) was observed in 42% of insertions and 56% of deletions at 30 °C, and in 58% of insertions and 71% of deletions at 35 °C. The detrimental effects were most pronounced at 35 °C and in residues located in the central region of the DHFR sequence, whereas residues in the N- and C-terminal regions were generally more tolerant to indels at 30 °C (**Fig. 2**). In general, many indels had similar effects in the WT, *ubr1*Δ and *san1*Δ strains, suggesting that the PQC E3s had minimal impact on the folding stability of these variants (**Fig. 2**). As expected, many central residues (positions 47–128) were particularly intolerant to indels across all three strains, indicating that indels at these positions disrupt protein folding and result in variants that remain unfolded or misfolded, even in the absence of Ubr1 or San1 (**Fig. 2**). However, some indel variants showed increased folding scores in the *ubr1*Δ and *san1*Δ strains compared to the WT strain, implying that more of these variants are stably folded in the absence of Ubr1 and San1 (**Fig. 3A**). Specifically, the proportion of well-folded insertions (CPOP folding score > 0.50) was 54% in the *ubr1*Δ strain and 51% in the *san1*Δ strain at 30 °C, compared with 47% in the WT strain. At 35 °C, the proportions were 39% in *ubr1*Δ and 40% in *san1*Δ, compared with 34% in the WT strain. The proportion of well-folded deletions (CPOP folding score > 0.50) was 45% in the *ubr1*Δ strain and 42% in the *san1*Δ strain at 30 °C, compared with 33% in the WT strain. At 35 °C, the proportions were 27% in *ubr1*Δ and 28% in *san1*Δ, compared with 23% in the WT strain. For both insertions and deletions, more variants were stabilized in the absence of Ubr1 at 30 °C, whereas Ubr1 and San1 displayed similar stabilizing effects at 35 °C (**Fig. 3A**). To visualize positions where indels could be rescued by deletion of the PQC E3s, we calculated the difference in folding scores between the WT strain and the *ubr1*Δ and *san1*Δ strains (**Fig. 3B**). This revealed numerous overlapping positions between insertions and deletions, most of which were located towards the N- and C-termini (**Fig. 3B**). Interestingly, the folding of many indels was rescued in both the absence of Ubr1 and San1, despite the distinct subcellular localizations and substrate recognition mechanisms of these E3 ligases (**Fig. 3B**). This highlights the overlapping roles of Ubr1 and San1 in recognizing and degrading misfolded DHFR indel variants to maintain proteostasis. Nevertheless, we observed differences in E3 specificity for certain indels, depending on temperature and their position within the protein (**Supplementary Fig. 4**). The degradation of indel variants positioned towards in the N- and C-terminal regions was mainly dependent on Ubr1 at 30 °C, whereas most indels in the N-terminal and central regions of DHFR were more strongly targeted by San1 at 35 °C (**Supplementary Fig. 4**).

**Fig. 2.**
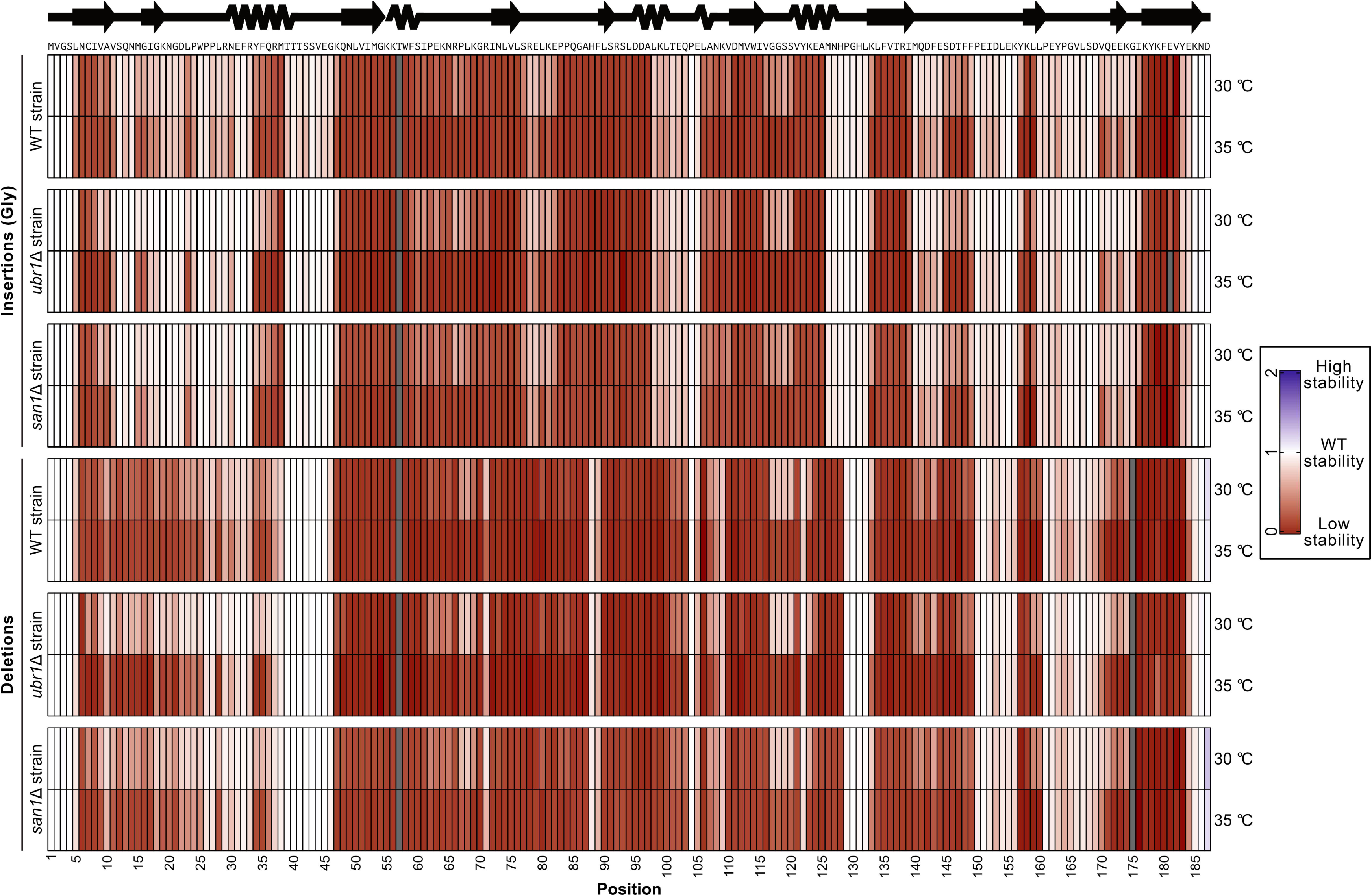
CPOP-based heatmaps of DHFR indel screen in WT and PQC E3 knockout strains. Heatmaps showing the folding and stability of DHFR insertions (top) and deletions (bottom) in the WT, *ubr1*Δ, and *san1*Δ strains at 30 °C and 35 °C. CPOP folding scores range from low stability (red), through wild-type-like stability (white), to high stability (blue). Missing variants are shown in grey. The amino acid sequence and a linear representation of human DHFR secondary structure are displayed above the heatmaps, and amino acid positions are shown below.

**Fig. 3.**
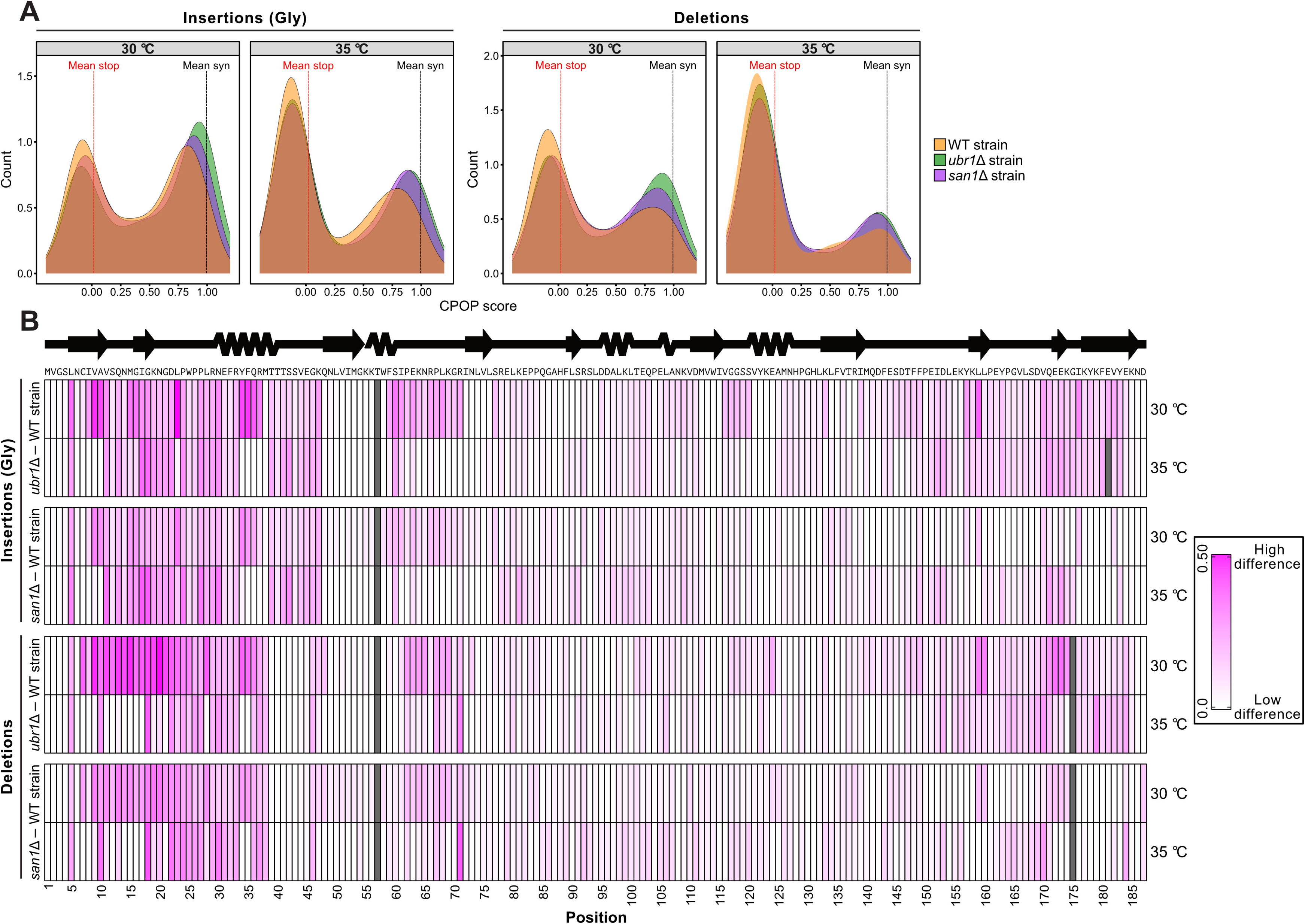
The effects of PQC E3 ligases on folding of the DHFR indel variants. (A) Score distributions of CPOP folding score for insertions (left) and deletions (right) in the WT (orange), *ubr1*Δ (green), and *san1*Δ (purple) strains at 30 °C and 35 °C. The CPOP folding scores were rescaled such that a score of 0 represents complete loss of stability (nonsense variants), while a score of 1 represents wild-type-like stability (synonymous variants). Scores greater than 1 indicate increased stability. The mean CPOP folding score for nonsense (red vertical line) and synonymous (black vertical line) variants were calculated across all tiles and strains. (B) Heatmaps showing the difference in folding and stability of DHFR insertions (top) and deletions (bottom) between the WT strain and the *ubr1*Δ and *san1*Δ strains at 30 °C and 35 °C. Differences were calculated by subtracting the CPOP folding score in the WT strain from the score in either the *ubr1*Δ or *san1*Δ strain. CPOP score differences range from low (white) to high (magenta). Missing variants are shown in grey. The amino acid sequence and a linear representation of human DHFR secondary structure are displayed above the heatmaps, and amino acid positions are shown below.

### The indel variants that are stabilized are partially functional and temperature-sensitive

Since we expected indel variants that are stabilized and well-folded in the absence of PQC E3 ligases to be only partially misfolded, we next asked whether these variants also exhibit temperature sensitivity (TS) and methotrexate (MTX) dependence. MTX is a small molecule that binds to the active site of DHFR, inhibits its enzymatic activity and stabilizes the native fold (16, 22, 24, 25). We previously assessed the folding of DHFR indel variants at 37 °C and in the presence of MTX to calculate a ΔTemperature score (score at 30 °C minus score at 37 °C) and a ΔMTX score (score with MTX minus score without MTX) for each variant (22). In this current study, we similarly calculated ΔUbr1 (score in *ubr1*Δ strain minus score in WT strain) and ΔSan1 (score in *san1*Δ strain minus score in WT strain) scores.

To identify indel variants most affected by temperature, MTX, Ubr1, or San1, we selected the top 25% of variants within each condition. This analysis revealed that the largest proportion of insertions (25.0%) and deletions (32.4%) were shared across all four conditions (TS, MTX, Ubr1, and San1) (**Fig. 4**), suggesting that these variants are hypomorphic (i.e., only marginally destabilized yet still recognized and targeted for proteasomal degradation by PQC E3 ligases). Importantly, these variants can be structurally rescued and stabilized through multiple mechanisms, including reduced temperature, ligand binding, or deletion of PQC E3 ligases, indicating that their native fold is not irreversibly disrupted.

**Fig. 4.**
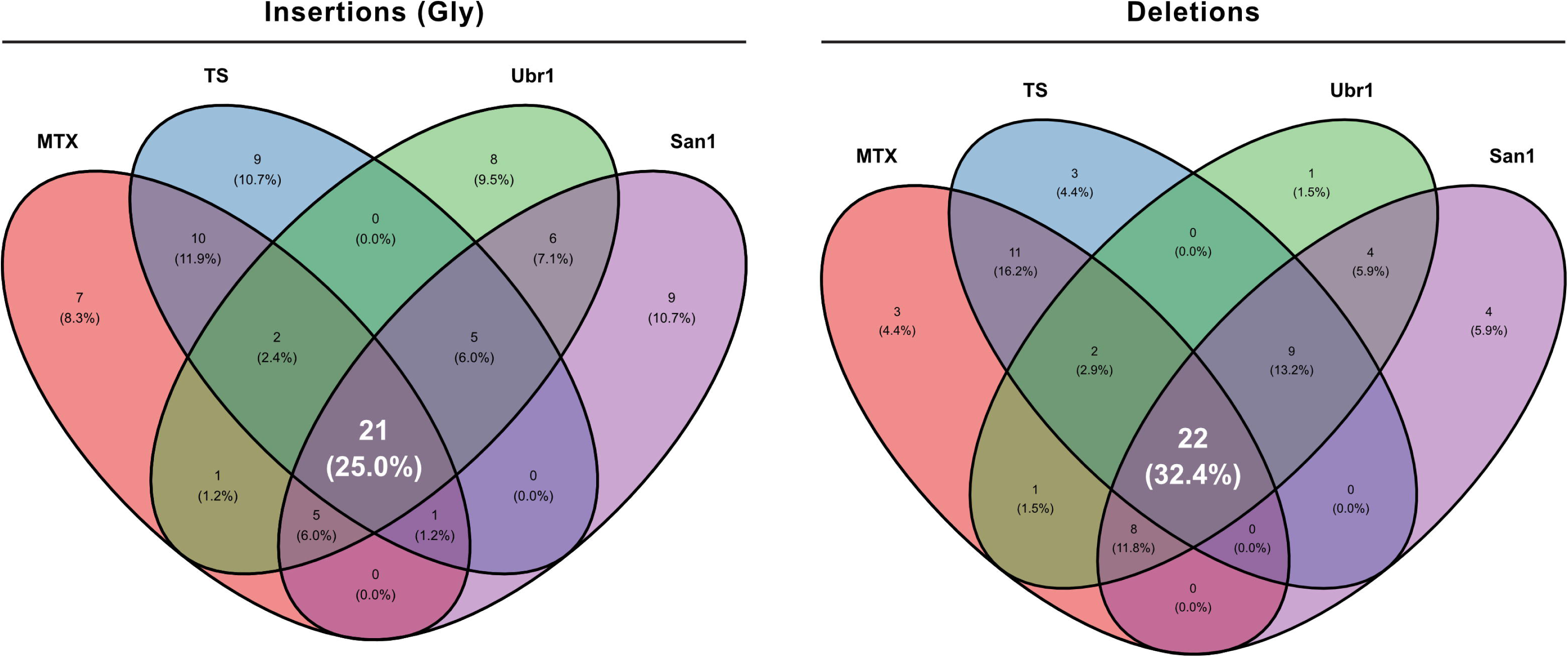
Phenotype distributions of DHFR indel variants. Venn diagrams showing the distribution of DHFR insertions (left) and deletions (right) across four categories: TS, MTX, Ubr1, and San1. Only the top 25% of indel variants that are most temperature-sensitive (TS), MTX-dependent (MTX), Ubr1-dependent (Ubr1), or San1-dependent (San1) are included. Numbers in each section represent the count of indels unique to or shared between conditions, with percentages shown in parentheses. The central overlapping regions indicate 21 insertions (25.0%) and 22 deletions (32.4%) common to all four conditions.

### Biophysical and cellular analysis of selected DHFR indels confirm partial folding defects

To further corroborate the results of the indel screen, we selected wild-type DHFR and nine indel variants for additional analysis. The variants were expressed as GFP fusions in wild-type cells at 30 °C and 35 °C and analysed by western blotting (**Fig. 5A**). Two variants, L74insG and L98Δ, displayed strongly reduced abundances at both temperatures, in line with their complete lack of folding in the CPOP-based screen (**Table 1**). The other selected indel variants appeared abundant at 30 °C but displayed reduced abundance at 35 °C (**Fig. 5A**), suggesting that several of these are temperature sensitive for folding, as was also evident from the screen (**Table 1**).

**Fig. 5.**
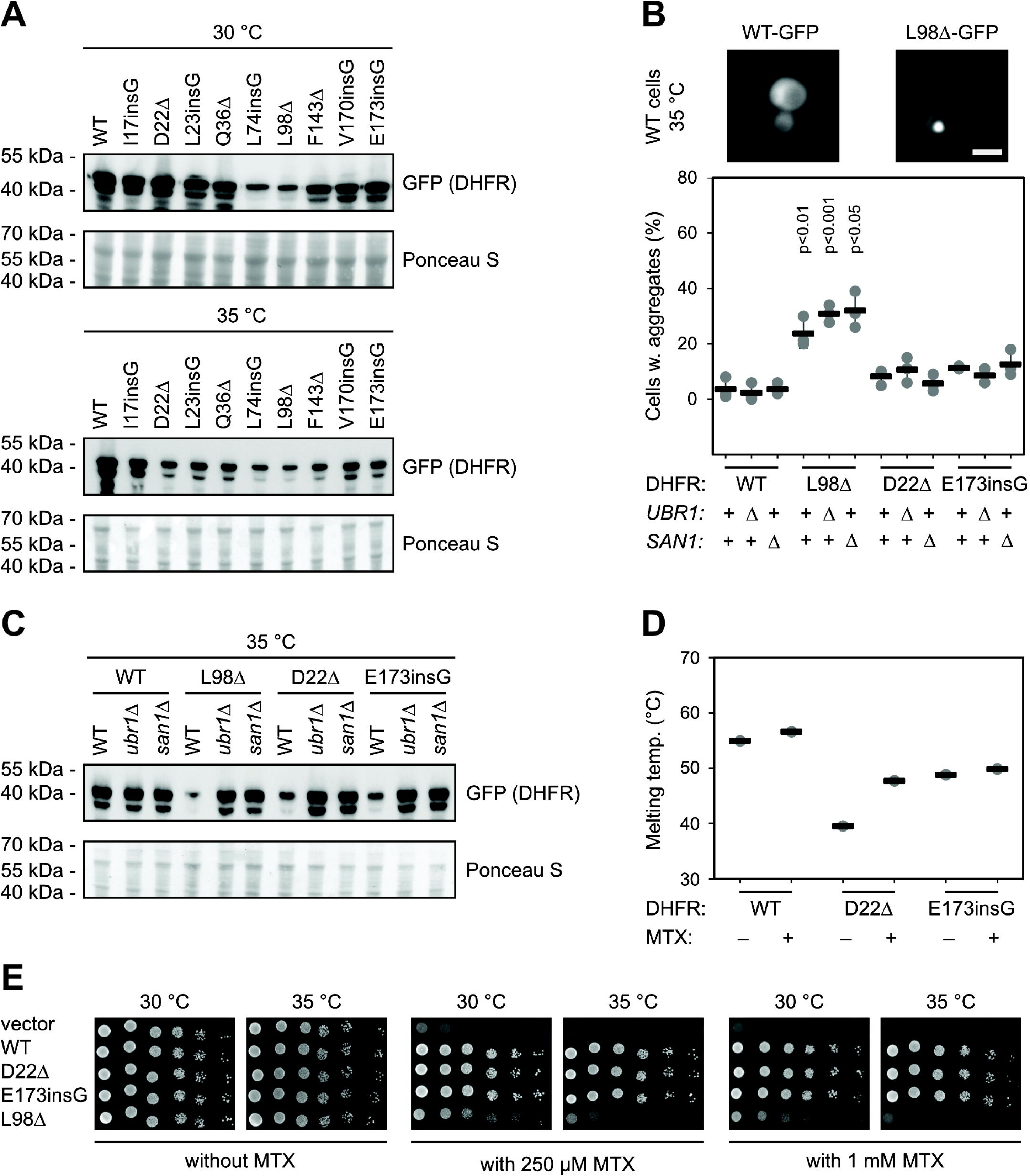
The indel variants display reduced abundance and structural stability. (A) The steady-state protein levels of the indicated DHFR indel variants fused to GFP were determined by SDS-PAGE and western blotting from wild-type yeast cells cultured at 30 °C (upper panel) and 35 °C (lower panel). Faster migrating bands were evident. Ponceau S staining was used as a loading control. (B) Live cell fluorescence microscopy of the cells expressing wild-type or L98Δ DHFR fused to GFP. Bar: 25 µm. Quantification of cells containing aggregates of selected DHFR variants and in wild-type cells or cells deleted (Δ) for UBR1 or SAN1 as indicated. Individual data points of three independent experiments are shown (points) along with the mean (black bars). The error bars indicate the standard deviation. Significance compared to WT DHFR was determined based on a t-test. The plotted data are included in the supplementary file. (C) The steady-state protein levels of the indicated DHFR indel variants fused to GFP were determined by SDS-PAGE and western blotting from wild-type (WT), *ubr1*Δ and *san1*Δ yeast cells cultured at 35 °C. Ponceau S staining was used as a loading control. (D) Melting points of purified DHFR variants determined by nanoDSF in the absence (-) or presence (+) of 50 µM methotrexate (MTX). Individual data points of three independent experiments are shown (points) along with the mean (black bars). The error bars indicate the standard deviation. The plotted data are included in the supplementary file. (E) Wild-type cells with vector, WT DHFR, D22Δ, E173insG and L98Δ fused to GFP were analyzed for growth on solid media at 30 °C and 35 °C, with 1 mM sulfanilamide and either 0, 250 µM or 1 mM MTX.

**Table 1.** Screening data for selected DHFR variants

|  | **WT** |  | ***ubr1*Δ** |  | ***san1*Δ** |  |  |
| --- | --- | --- | --- | --- | --- | --- | --- |
| **Variant** | **30 °C** | **35 °C** | **30 °C** | **35 °C** | **30 °C** | **35 °C** | **Interpretation\*** |
| I17SinsG | 0.677 | 0.319 | 0.962 | 0.688 | 0.890 | 0.712 | Moderate ts |
| D22Δ | 0.430 | 0.000 | 0.865 | 0.327 | 0.756 | 0.390 | Strong ts |
| L23insG | 0.290 | -0.042 | 0.773 | -0.002 | 0.637 | 0.122 | Strong ts |
| Q36Δ | 0.457 | 0.010 | 0.800 | 0.187 | 0.690 | 0.249 | Strong ts |
| L74insG | -0.096 | -0.118 | -0.108 | -0.131 | -0.045 | -0.172 | No folding |
| L98Δ | -0.140 | -0.183 | -0.091 | -0.116 | -0.087 | -0.097 | No folding |
| F143Δ | 0.470 | -0.081 | 0.580 | -0.086 | 0.477 | 0.026 | Strong ts |
| V170insG | 0.580 | -0.089 | 0.826 | 0.100 | 0.706 | -0.008 | Strong ts |
| E173insG | 0.709 | 0.108 | 0.877 | 0.344 | 0.763 | 0.364 | Strong ts |

\*: ts, temperature sensitive.

To probe the indel effects further, we studied the localization of wild-type DHFR, the severe L98Δ variant, and the intermediate D22Δ and E173insG variants by live-cell fluorescence microscopy. As expected, the variants localized to the cytosol, however, for about 30% of the cells the severe L98Δ variant was localized as a large aggregate (**Fig. 5B**). This was not evident for the intermediate indel variants and was not further aggravated in the E3 deletion strains (**Fig. 5B**). However, as assessed by western blotting, at 35 °C the abundance of the indel variants increased in the E3 deletions (**Fig. 5C**).

Next, we tested if the variants are indeed structurally destabilized. We produced the D22Δ and E173insG protein variants along with wild-type DHFR as 6His-SUMO fusions in *E. coli* and purified the proteins (**Supplementary Fig. 5**). Given the severe nature of the L98Δ variant, we did not attempt to purify this protein. After cleavage of the 6His-SUMO fusions, the thermal unfolding of the DHFR variants was determined by nano Differential Scanning Fluorimetry (nanoDSF) (**Fig. 5D**). As expected, the indel variants displayed strongly reduced melting temperatures (39.5 °C for D22Δ and 48.8 °C for E173insG, compared with 54.9 °C for wild-type DHFR). In the presence of MTX the melting temperatures increased to 47.7 °C for D22Δ and 49.9 °C for E173insG and 56.6 °C for wild-type DHFR (**Fig. 5D**).

Using the CPOP system we confirmed the temperature sensitive folding of the D22Δ and E173insG variants and their increased folding (growth) in the *san1*Δ and *ubr1*Δ backgrounds (**Supplementary Fig. 6)**. Finally, we tested if the selected indel variants were able to confer resistance to MTX in yeast cells when overexpressed. Indeed, this appeared to be the case for D22Δ and E173insG, while expression of the severe L98Δ variant only led to minor MTX resistance (**Fig. 5E**). Based on these results, we conclude that D22Δ and E173insG, are only partially unfolded and potentially functional hypomorphic variants. We speculate that many of the other indel variants that are targeted by Ubr1 and San1 behave similarly to D22Δ and E173insG.

## Discussion

Most proteins fold into stable structures, but stress conditions such as heat, mutations, or missing cofactors can lead to loss of structural stability, misfolding and harmful non-native interactions with other cellular components. To manage this, cells rely on the PQC system, where molecular chaperones help refold misfolded proteins, while E3 ubiquitin ligases, like yeast Ubr1 and San1, target them for proteasomal degradation. This degradation prevents toxic protein build-up and maintains proteostasis but can also lead to the turnover of partially unfolded but still functional proteins, potentially impairing vital functions (3, 5, 26).

Using a yeast-based protein folding reporter, we examined how indel mutations affect folding of the DHFR protein in strains lacking Ubr1 and San1. We found that many indels, especially those with mutations near the protein’s termini, were stabilized in the E3 knockouts. We found that both Ubr1 and San1 were able to target several indels for degradation, consistent with previous studies showing that the two E3s function in parallel (8, 27–31). It appeared that the degradation was mostly Ubr1-dependent at 30 °C, but at 35 °C this effect was less pronounced, and some indels became better targets of San1. This may reflect increased cellular stress at elevated temperatures, which requires dual action of Ubr1 and San1 to maintain proteostasis. Alternatively, it could result from increased protein unfolding, which exposes hydrophobic regions and thereby enhances substrate recognition by San1 (12). Many of the variants that were stabilized in the E3 knockouts were also stabilized by lower temperatures or methotrexate binding and were primarily positioned towards the N- and C-terminal regions of DHFR. This agrees with other systematic studies (32–39), that suggest indels are better tolerated in flexible loops and at protein termini, presumably because these regions are less critical for maintaining the stability of the protein core. In agreement with our previous analyses (22), we observed that insertions are generally better tolerated than deletions, likely because proteins can more easily accommodate extra residues, such as by extending loops, whereas deletions may disrupt structure by introducing strain or removing critical elements.

Although the structural stability of a protein, in general, correlates with the abundance of the protein in the cell (40, 41), this correlation can collapse, e.g. when mutations in exposed or disordered regions of a protein generate a degradation signal (degron) (42) or if the PQC system is compromised (17, 43). When proteasomal degradation is blocked by E3 knockouts, a larger amount of the indel protein is retained in the native state or remains unfolded, possibly leading to aggregation (**Fig. 6**). Accordingly, a protein folding sensor such as CPOP, which directly measures a protein’s folding state within the cellular environment, may provide more accurate insights into protein stability than abundance sensors like GFP fusions, which only report on protein levels and cannot distinguish folded from unfolded or aggregated conformations. Thus, although we observed an increased abundance of L98Δ fused to GFP in the E3 mutants, the CPOP results indicate that this variant does not form a stable fold, so the increased abundance likely reflects aggregation of the misfolded protein. However, the folding reporter comes with its own limitations. For instance, certain indels in the context of the CPOP folding sensor may be degraded more efficiently due to the concomitant unfolding of CPOP, which might be further aggravated since CPOP is a dimer (23) and therefore likely unfolds in a cooperative manner. In theory this should make the CPOP system more sensitive than e.g. a GFP fusion. However, for the variants we analyzed in low throughput, the CPOP folding in general correlated with the protein abundance of the GFP fused indels.

**Fig. 6.**
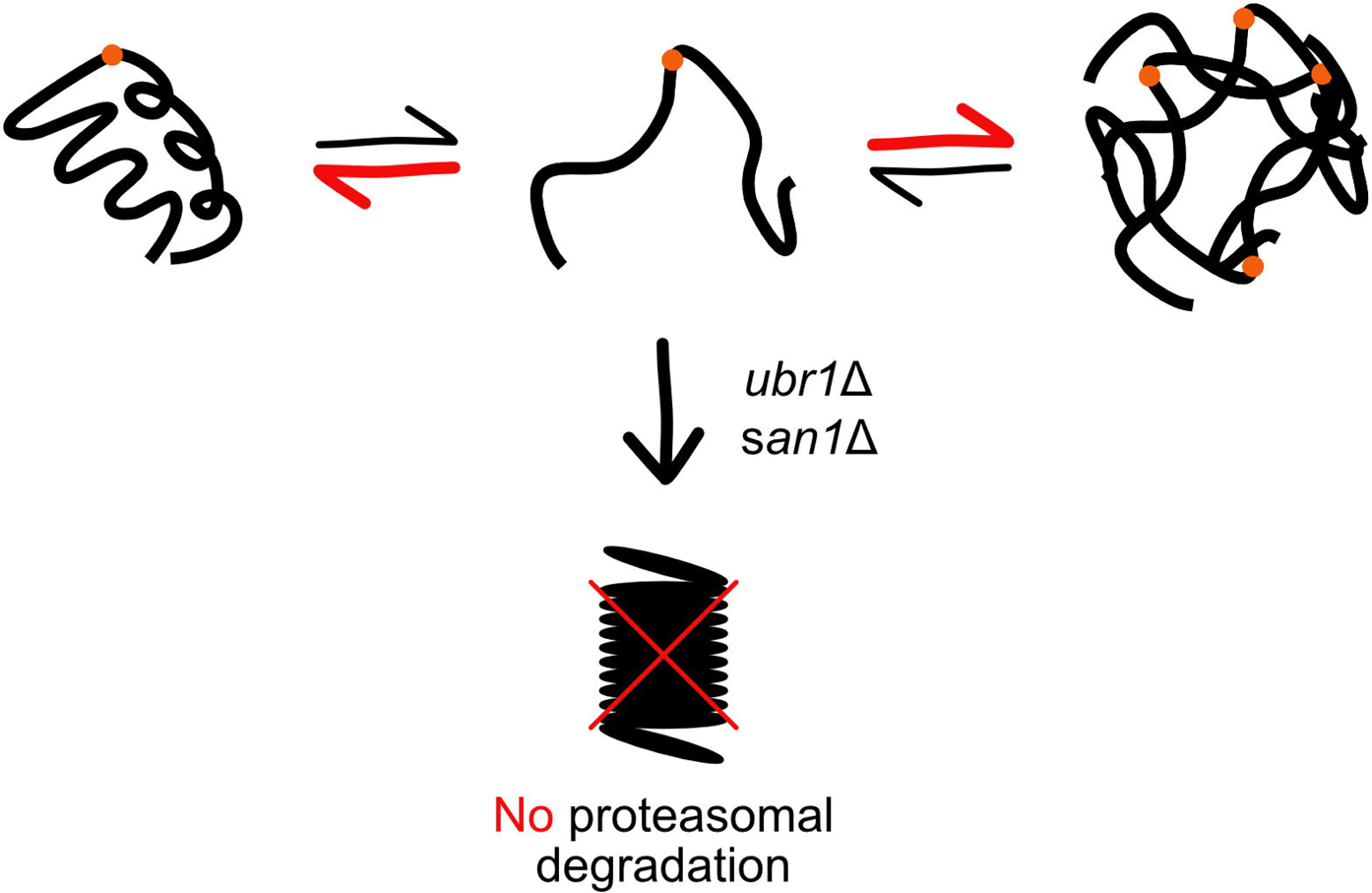
Model of indel PQC degradation. The presented data are consistent with a model in which insertions and deletions (red dot) in DHFR lead to unfolding of the protein. The unfolded DHFR indel variants are targeted for proteasomal degradation via the E3s Ubr1 and San1. In E3 deletion strains, proteasomal degradation is reduced and an increased amount of the protein is found in either the native state or as an aggregate.

Studying the folding and degradation of indel proteins can aid in developing therapies for genetic diseases by rescuing partially functional proteins and improving variant interpretation. In biotechnology, it can guide protein engineering and enable better control over protein stability and function.

## Methods

### Plasmids and cloning

Codon-optimized DNA sequences for yeast expression of the empty CPOP construct and wild-type *E. coli* OPRTase (UniProt ID: P0A7E3) were synthesized and cloned into the pDONR221 vector by Genscript. All human DHFR (UniProt ID: P00374) indel variants for small scale testing were purchased from Genscript, either inserted into the CPOP construct or C-terminally fused to GFP. For expression in yeast cells, entry clones were Gateway-cloned (Invitrogen) into pAG415GPD-ccdB (Addgene plasmid #14146; http://n2t.net/addgene:14146; RRID:Addgene_14146). Codon-optimized DNA sequences of wild-type DHFR and indel variants for *E. coli* expression were synthesized, fused to 6His-SUMO1, and cloned into the pET28a vector by Genscript.

### Yeast strains and media

The *ura5*Δ*ura10*Δ yeast strain has been described before (22). The *ubr1*Δ and *san1*Δ yeast strains were obtained from the Euroscarf collection. The *ura5*Δ*ura10*Δ*ubr1*Δ and *ura5*Δ*ura10*Δ*san1*Δ yeast strains were constructed by homologous recombination of PCR products using long oligos (**Supplementary File 1**) on a hygromycin resistance cassette. Correct integration was confirmed by PCR using oligos flanking the regions of homology (**Supplementary File 1**).

Small scale yeast transformations were performed as described in (44). Yeast cells were cultured in yeast extract, peptone, dextrose (YPD) medium (2% (w/v) glucose, 2% (w/v) tryptone, 1% (w/v) yeast extract), and synthetic complete (SC) medium (2% (w/v) glucose, 0.67% (w/v) yeast nitrogen base without amino acids (Sigma), 0.2% (w/v) synthetic drop-out supplement (Sigma)). Solid SC media was prepared using synthetic drop-out supplement (Sigma).

### Library design, transformation and selection

The DHFR indel libraries were designed as previously described in (22). The libraries were transformed in parallel into the *ura5*Δ*ura10*Δ strain, the *ura5*Δ*ura10*Δ*ubr1*Δ strain, and the *ura5*Δ*ura10*Δ*san1*Δ as previously described in (22).

Once the transformation cultures reached saturation, triplicate samples corresponding to 9 OD_600nm_ units were collected by centrifugation (17,000 g, 1 minute, room temperature) and the resulting cell pellets were stored at -20 °C. These samples represented the pre-selection control. For selection, triplicates of 2×5 OD_600nm_ units were collected, washed three times with sterile water (17,000 g, 1 minute, room temperature), and resuspended in 400 μL sterile water. Subsequently, 100 μL of each suspension was plated onto square selection plates (144 cm^2^). Plates were briefly air-dried and incubated at 30 °C or 35 °C for two days. Following incubation, confluent lawns of yeast colonies were present on all plates. Cells were scraped into 10 mL of sterile water. Then 9 OD_600nm_ units were harvested per plate (17,000 g, 1 minute, room temperature) and the pellets were stored at -20 °C. These samples represented the post-selection conditions.

### Amplicon preparation and library sequencing

Plasmid DNA was extracted from the collected cell pellets using the ChargeSwitch Plasmid Yeast Mini kit (Invitrogen), and DNA concentrations were quantified using the Qubit 2.0 Fluorometer (Invitrogen) along with the Qubit dsDNA HS Assay Kit (Thermo Fisher Scientific). Amplicons were prepared and sequenced as previously described (22).

### Analysis of sequencing data

The resulting paired-end sequencing reads were processed as previously described (22). Briefly, merged and trimmed reads were processed to count unique DNA sequences, which were then used as input for our in-house Python script (22). This script assigned each sequence a single mutation type (insertion, deletion, synonymous, or nonsense) and a corresponding position (1–187). As an additional filtering step, variants with fewer than 10 reads in the pre-selection condition were excluded to reduce sequencing noise. Finally, a rescaled score was computed for each mutation type at 30 °C and 35 °C by normalizing to synonymous wild-type variants, where a score of 0 represents the average of nonsense variants and a score of 1 represents the average of synonymous variants. This resulted in six scores for each indel variant in DHFR: 1) *ura5*Δ*ura10*Δ strain (referred to as WT strain) at 30°C, 2) *ura5*Δ*ura10*Δ strain (referred to as WT strain) at 35 °C, 3) *ura5*Δ*ura10*Δ*ubr1*Δ strain at 30 °C, 4) *ura5*Δ*ura10*Δ*ubr1*Δ strain at 35 °C, 5) *ura5*Δ*ura10*Δ*san1*Δ strain at 30 °C, and 6) *ura5*Δ*ura10*Δ*san1*Δ strain at 35 °C. For technical reasons related to sequencing, the following replicates were excluded: *ura5*Δ*ura10*Δ*ubr1*Δ strain, tile 2, replicate 3, 35 °C; *ura5*Δ*ura10*Δ*ubr1*Δ strain, tile 3, replicate 3, 35 °C.

### Electrophoresis and blotting

Whole cell extracts from yeast were prepared using NaOH as previously described (45). For SDS-PAGE the samples were resolved on 12.5% acrylamide gels. For western blotting the proteins were transferred to 0.2 μm nitrocellulose membranes. Staining with Ponceau (0.1 % (w/v) Ponceau S in 5% (v/v) acetic acid) was used as a loading control. The membranes were blocked in PBS (10 mM Na_2_HPO_4_, 1.8 mM KH_2_PO_4_, 137 mM NaCl, 3 mM KCl, pH 7.4) containing 5% (w/v) milk powder and 5 mM NaN_3_. The primary antibody (diluted 1:1000) was rat anti-GFP monoclonal (clone 3H9) (Chromotek, 3H9). The secondary antibody (diluted 1:5000) was horse radish peroxidase-conjugated goat anti-rat IgG (Proteintech, SA0001-15). Uncropped western blots are provided in the supplementary material (**Supplementary File 1**). The blots were developed using the ECL detection reagent (Amersham GE Healthcare) and a ChemiDoc MP Imaging System (Bio-Rad). Coomassie staining of SDS-PAGE gels was performed using 2.5 g/L Coomassie Brilliant Blue R250 in 45% (v/v) ethanol and 10% (v/v) acetic acid.

### Microscopy

For microscopy, live yeast cells in exponential phase were washed in PBS and directly imaged using a Zeiss Axio Imager Z1 microscope attached to a Hamamatsu ORCA-ER digital camera.

### Yeast growth assays

Yeast cultures for growth assays were incubated overnight (30 °C) until reaching exponential phase. Cells were then collected by centrifugation (3,000 g, 5 minutes), washed twice with sterile water, and resuspended in sterile water. Cell suspensions were normalized to an OD_600nm_ of 0.4 and subjected to five-fold serial dilutions in sterile water. From each dilution, 5 μL was spotted onto agar plates. The plates were allowed to air dry before incubation at the indicated temperature for 2-3 days.

### Protein purification

Bacterial cultures in exponential phase were grown in LB media supplemented with 2 mM folic acid and 100 µg/mL ampicillin, were induced with 0.1 M isopropyl β-D-1-thiogalactopyranoside (IPTG) at 16 °C for 16-18 hours. Cells were harvested by centrifugation (3,000 g, 20 min) and the pellet resuspended in 40 mL lysis buffer (50 mM NaH_2_PO_4_, 300 mM NaCl, 1 mM phenylmethylsulfonyl fluoride (PMSF), Complete protease inhibitors without EDTA (Roche), pH 7.4). The cells were lysed by sonication on ice, and the lysates were cleared by centrifugation (13,000 g, 30 min., 4 °C). The cleared lysate was incubated with 1 mL TALON metal affinity resin (Clontech) at 4 °C for 1 hour. Unbound protein was washed out with 30 column volumes (CVs) of wash buffer (50 mM NaH_2_PO_4_, 300 mM NaCl, 10 mM imidazole, pH 7.4), and the bound protein was eluted with 5 CVs elution buffer (50 mM NaH_2_PO_4_, 300 mM NaCl, 250 mM imidazole, pH 7.4). Samples of flow-through, wash and elution were separated by SDS-PAGE followed by Coomassie staining to determine which fractions contained the fusion protein (**Supplementary Fig. 5**). Samples containing the fusion protein were pooled and dialyzed against cleavage buffer (25 mM Na-phosphate, 510 mM NaCl, 0.5 mM folate, 0.5 mM DTT, pH 7.4) at 4 °C overnight. The fusion protein was cleaved with 1:100 dilution of Ulp1 6His-SUMO protease for 2 hours at 4 °C. The samples were incubated with 1 mL TALON metal affinity resin (Clontech) for 30 minutes at 4 °C. Cleaved protein was washed out with 10 CVs lysis buffer and bound protein was eluted with 5 CVs elution buffer. Fractions were analyzed by SDS-PAGE and Coomassie staining (**Supplementary Fig. 5**). Fractions containing cleaved DHFR were pooled and dialyzed against storage buffer (50 mM K-phosphate, 100 mM KCl, 1 mM DTT, 25 μM folic acid, pH 6.5) at 4 °C overnight. The purified protein was stored at 4 °C for no longer than 2 days before analyses by nanoDSF.

### Heat denaturation

Purified samples of each of the DHFR protein variants in storage buffer (see above) were used for heat denaturation. To replace bound folate, 50 μM MTX was added to the samples followed by 5 min incubation at RT. Then, 15 μL samples were loaded by placing a high sensitivity quartz capillary (Nanotemper) in the samples. The unfolding was followed by intrinsic tryptophan fluorescence at 350 nm after excitation at 280 nm using a Nanotemper Prometheus NT.48 fluorimeter. The temperature was ramped with 1 °C/min from 20 °C to 95°C. The numerical data are included in the supplementary material (**Supplementary File 1**).

### Quantification and statistical analysis

The screens were performed in triplicates. Data are displayed as individual points with the mean and standard deviation indicated. For the aggregate quantification, the number of cells containing aggregates from manual counting of at least 100 cells in three independent experiments and standard deviations are presented. Significance was determined based on a t-test. The numerical data are included in the supplementary material (**Supplementary File 1**).

## Acknowledgements

We acknowledge the use of the sequencing and computing core facilities at the Department of Biology, University of Copenhagen. We thank Anne-Marie Lauridsen for technical assistance. Parts of Fig. 1 were created with BioRender.com.

## Competing interests

The authors have no relevant financial or non-financial interests to disclose.

This article includes the following supplementary information.

## Supplemental files

- Supplementary figures and tables (SupplementaryMaterial.pdf).
- Supplementary dataset (SupplementaryFile.xlsx).

## Data and code availability

All data generated are included in the figures and supplementary files. Sequencing reads are available at the NCBI Gene Expression Omnibus (GEO) repository (accession number: GSE302176). The code is available at GitHub (https://github.com/KULL-Centre/_2025_Larsen-Ledet_CPOP2).

## Author contributions

S.L.-L., C.H.S., C.M.H. and S.G. performed the experiments. S.L.-L., C.H.S., S.G., A.S. and K.T. analyzed the data. S.L.-L., S.G. and R.H.-P. conceived the study. S.L.-L. prepared the figures. S.L.-L. and R.H.-P. wrote the paper.

