## Supplementary Material for "Blocking protein quality control degradation leads to structural stabilization of DHFR indel variants"

Sven Larsen-Ledet[1,#], Caroline H. Suhr[1,#], Sarah Gersing[1], Celeste M. Hackney[1], Amelie Stein[1], Kaare Teilum[1], and Rasmus Hartmann-Petersen[1,*]

1: Department of Biology, University of Copenhagen, Ole Maaløes Vej 5, DK-2200 Copenhagen, Denmark.

#: These authors contributed equally.

**Supplementary Fig. 1** – *Correlations between repeats for wild-type cells.* p.2
**Supplementary Fig. 2** – *Correlations between repeats for san1Δ cells.* p.3
**Supplementary Fig. 3** – *Correlations between repeats for ubr1Δ cells.* p.4
**Supplementary Fig. 4** – *San1- and Ubr1-specific sites.* p.5
**Supplementary Fig. 5** – *Purification of DHFR variants from E. coli.* p.6
**Supplementary Fig. 6** – *Growth assays for selected variants.* p.7

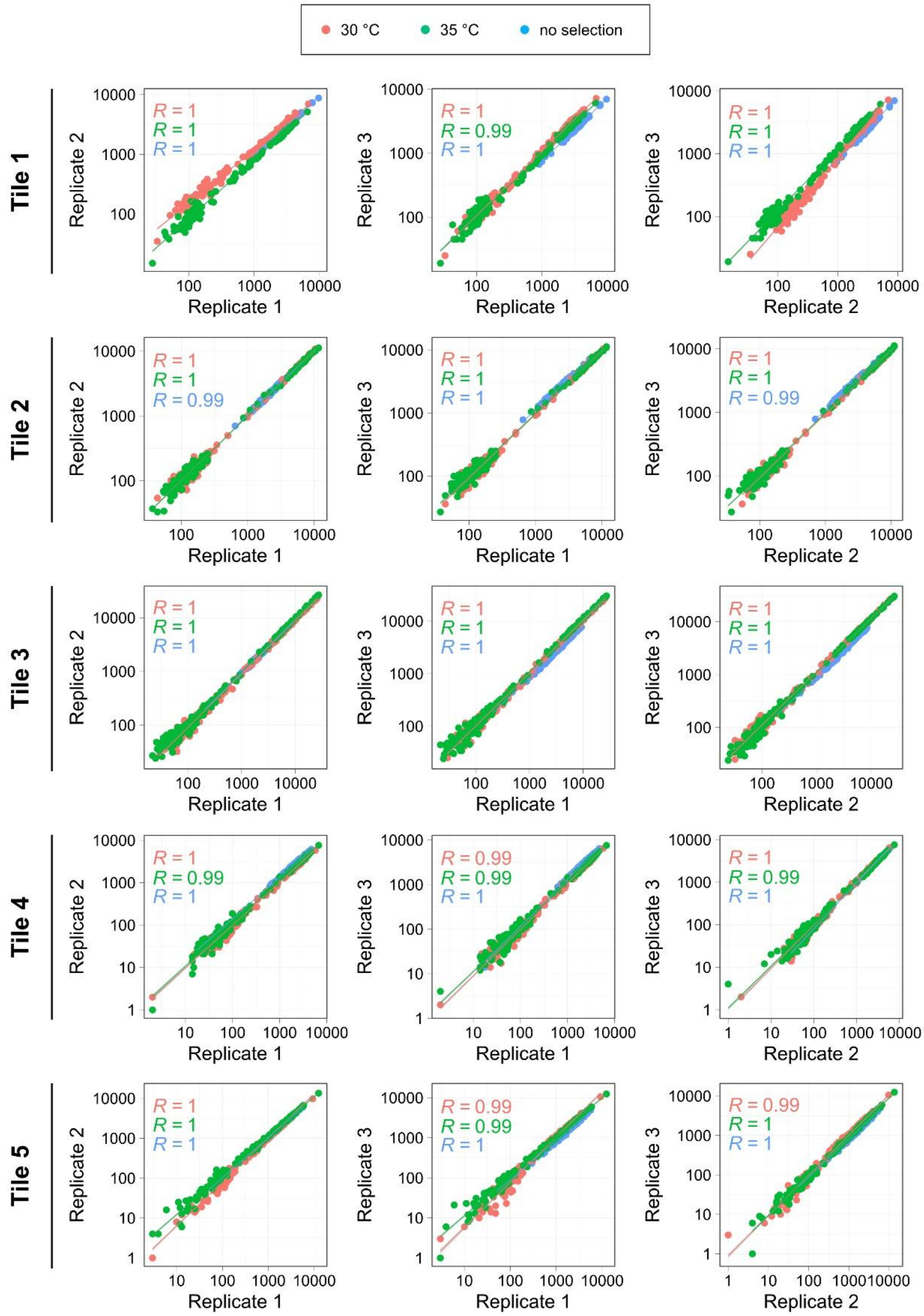


**Supplementary Fig. 1** – *Correlations between repeats for wild-type cells.*
Correlation plots of the three repeats (replicate 1-3) for the five tiles (tile 1-5) of DHFR at the three conditions: 30 °C (red), 35 °C (green), and no selection (control, blue).

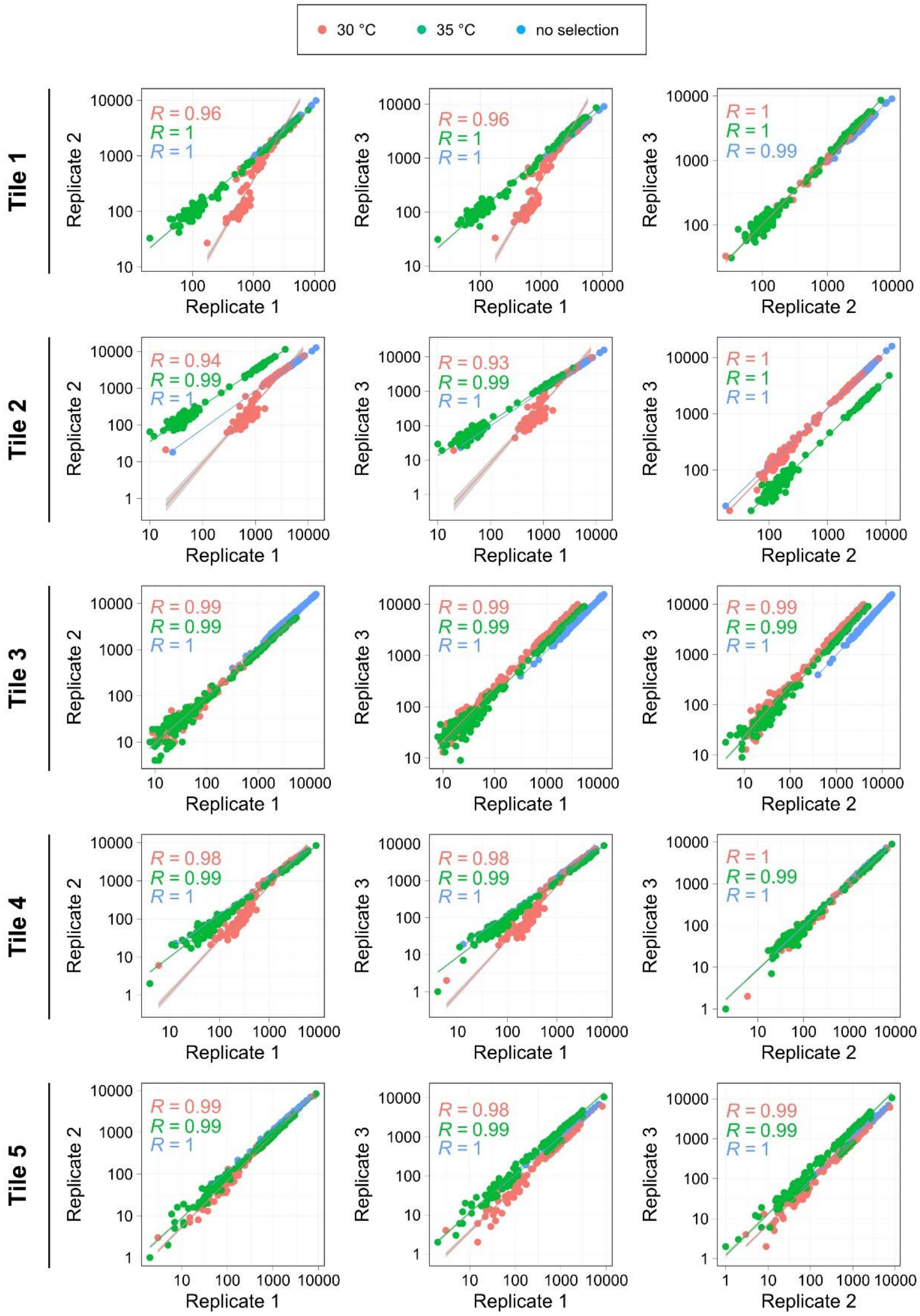


**Supplementary Fig. 2** – *Correlations between repeats for san1Δ cells.*
Correlation plots of the three repeats (replicate 1-3) for the five tiles (tile 1-5) of DHFR at the three conditions: 30 °C (red), 35 °C (green), and no selection (control, blue).

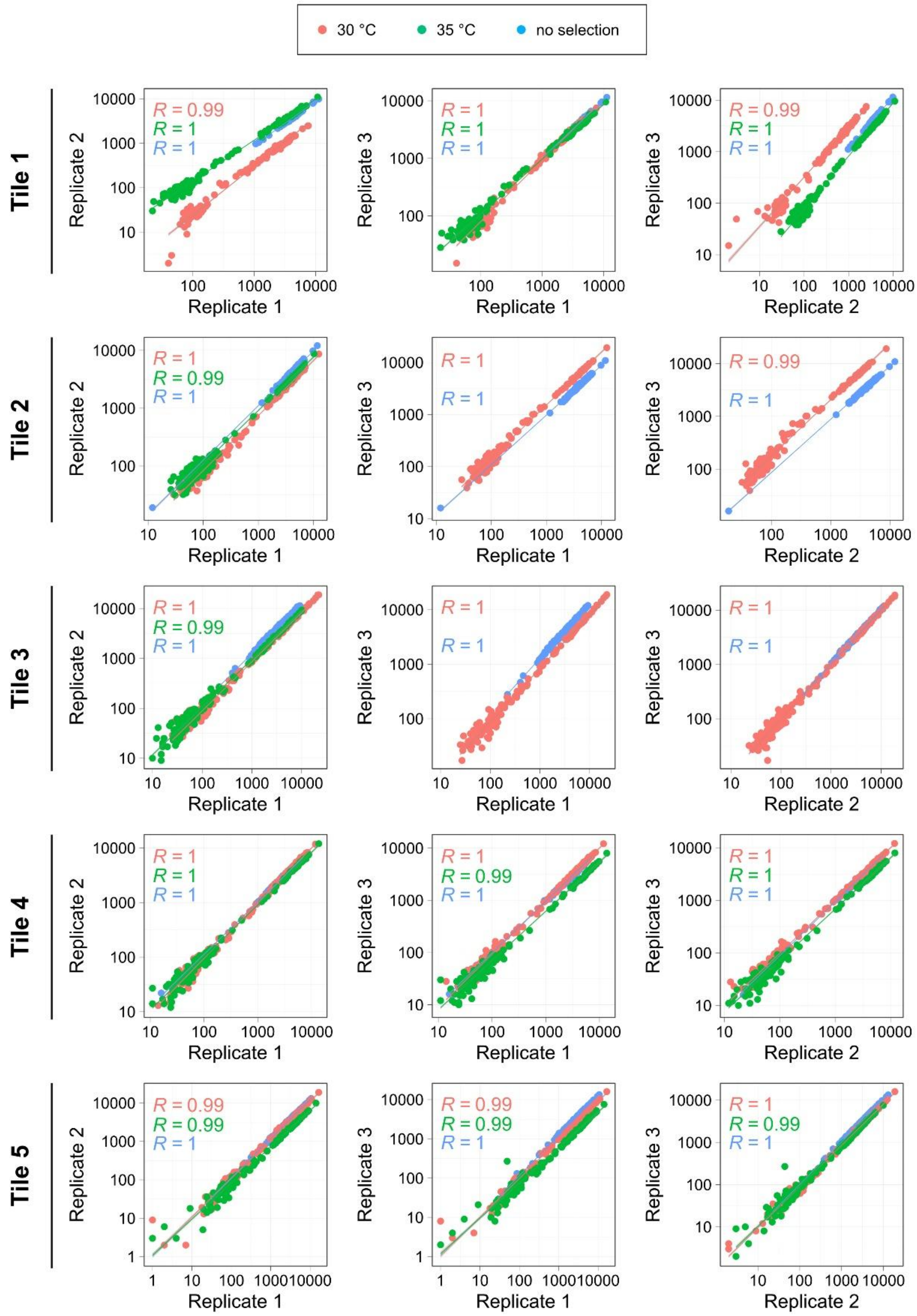


**Supplementary Fig. 3** – *Correlations between repeats for ubr1Δ cells.*
Correlation plots of the three repeats (replicate 1-3) for the five tiles (tile 1-5) of DHFR at the three conditions: 30 °C (red), 35 °C (green), and no selection (control, blue).

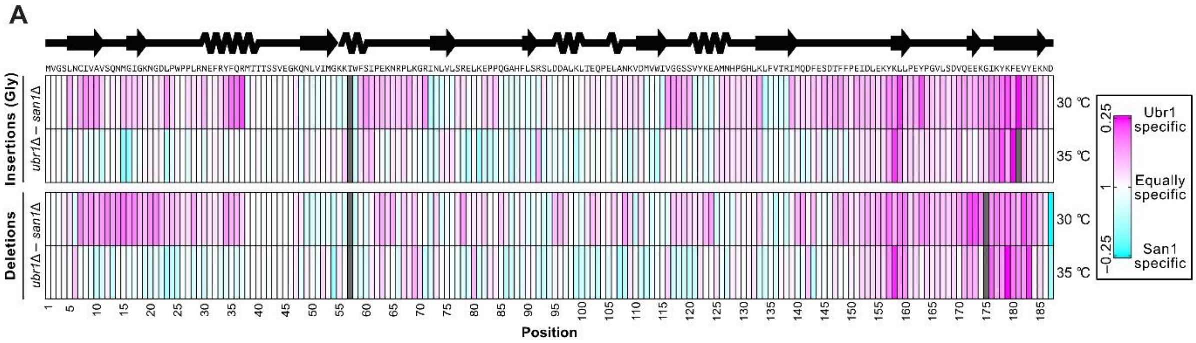


**Supplementary Fig. 4** – *San1- and Ubr1-specific sites.*

Heat-map of the difference in the *ubr1Δ* and *san1Δ* scores. The position of the indel in DHFR is on the x-axis. San1-specific sites are shown in cyan- Ubr1-specific sites are shown in pink.

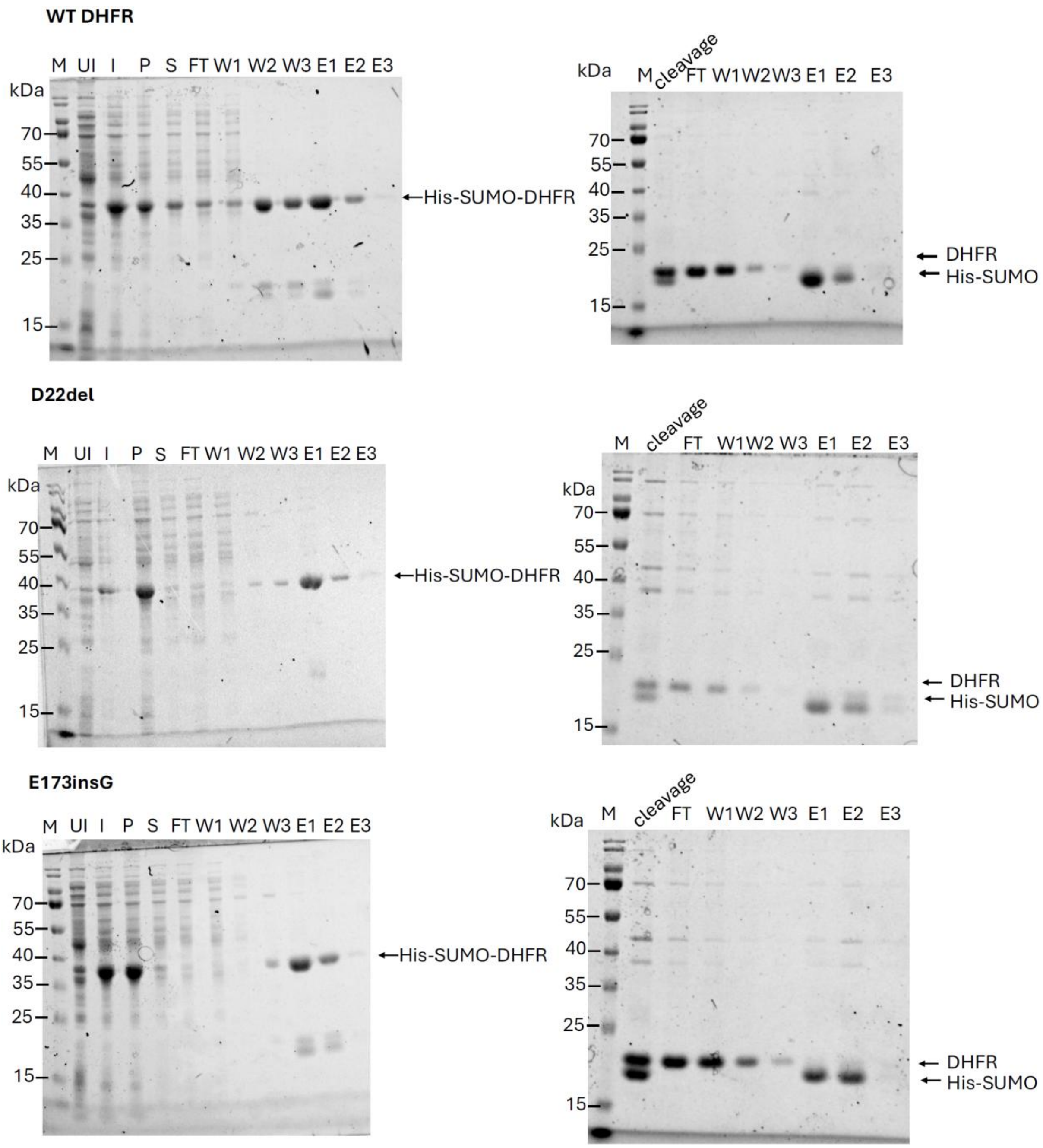


**Supplementary Fig. 5** – *Purification of DHFR variants from E. coli.*
DHFR wild-type (WT) (upper panels), D22Δ (central panels) and E173insG (lower panels), were expressed in *E. coli* fused to 6His-SUMO (uninduced, UI and induced, I) and separated by centrifugation into an insoluble (pellet, P) and soluble (supernatant, S) fraction. The soluble fraction was purified by passing cleared lysate over a TALON resin (flow through), washing (wash, W1, W2 and W3), and eluting with imidazole (eluate, E1, E2 and E3). The eluted fractions E1 and E2 were pooled and cleaved with 6His-tagged Ulp1 SUMO protease (cleavage). The cleaved material was passed over another TALON column, and the flow through (FT) and first wash fraction (W1) were collected. Subsequently, the His-SUMO fusion was eluted with imidazole (E1-E3). The samples along with a molecular weight marker (M) were resolved by SDS-PAGE and the gels were stained with Coomassie Brilliant Blue.

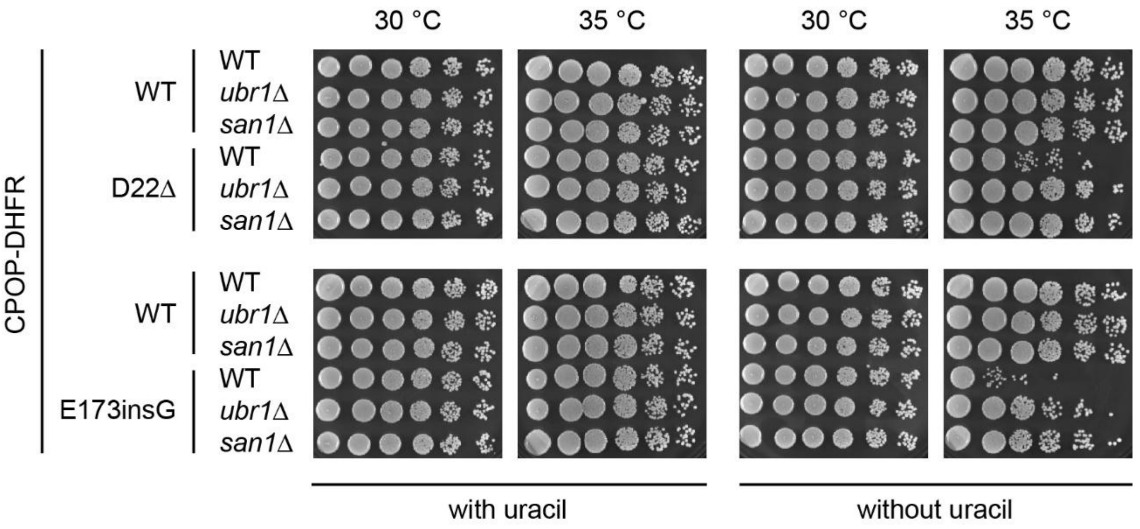


**Supplementary Fig. 6** – *Growth assays for selected variants.*
DHFR wild-type (WT), D22Δ and E173insG in the CPOP context were expressed in *ura5*Δ*ura10*Δ cells (WT) or *ura5*Δ*ura10*Δ cells deleted for the E3s Ubr1 (*ubr1*Δ) or San1 (*san1*Δ) as indicated. Cells in exponential phase were 5-fold serially diluted and spotted onto media with (left) or without (right) uracil and incubated 30 °C or 35 °C. Note that the indels display a temperature sensitive growth defect without uracil which is partially rescued by deletion of the E3s.
